# Ultrastructural analysis of prostate cancer tissue provides insights into androgen-dependent adaptations to membrane contact site establishment

**DOI:** 10.1101/2023.05.10.540238

**Authors:** Lisa M. Butler, Emma Evergren

## Abstract

Membrane trafficking and organelle contact sites are important for regulating cell metabolism and survival; processes often deregulated in cancer. Prostate cancer is the second leading cause of cancer-related death in men in the developed world. While early-stage disease is curable by surgery or radiotherapy there is an unmet need to identify prognostic biomarkers, markers to treatment response and new therapeutic targets in intermediate-late stage disease. This study explored the morphology of organelles and membrane contact sites in tumor tissue from normal, low and intermediate histological grade groups. The morphology of organelles in secretory prostate epithelial cells; including Golgi apparatus, ER, lysosomes; was similar in prostate tissue samples across a range of Gleason scores. Mitochondrial morphology was not dramatically altered, but the number of membrane contacts with the ER notably increased with disease progression. A three-fold increase of tight mitochondria-ER membrane contact sites was observed in the intermediate Gleason score group compared to normal tissue. To investigate whether these changes were concurrent with an increased androgen signaling in the tissue, we investigated whether an anti-androgen used in the clinic to treat advanced prostate cancer (enzalutamide) could reverse the phenotype. Patient-derived explant tissues with an intermediate Gleason score were cultured *ex vivo* in the presence or absence of enzalutamide and the number of ER-mitochondria contacts were quantified for each matched pair of tissues. Enzalutamide treated tissue showed a significant reduction in the number and length of mitochondria-ER contact sites, suggesting a novel androgen-dependent regulation of these membrane contact sites. This study provides evidence for the first time that prostate epithelial cells undergo adaptations in membrane contact sites between mitochondria and the ER during prostate cancer progression. These adaptations are androgen-dependent and provide evidence for a novel hormone-regulated mechanism that support establishment and extension of MAMs. Future studies will determine whether these changes are required to maintain pro-proliferative signaling and metabolic changes that support prostate cancer cell viability.

## Introduction

Prostate cancer (PCa) is the second most common cancer in men in the world. It typically begins as a slow growing androgen-dependent cancer before acquiring driver mutations which lead to increased proliferation, aggressiveness and metastatic potential. Prostate epithelial cells are specialized in producing and releasing proteins, metabolites and membrane vesicles which contribute to the production of seminal fluid. The molecular events and mutations that contribute to the transformation of prostate cells are increasingly well defined with a range of functional studies implicating them in metabolic dysregulation, cell proliferation and cell survival. The impact of these changes on cellular ultrastructure as defined by the morphology, distribution and relationship between organelles such as mitochondria and the endoplasmic reticulum is by contrast largely uncharacterized. These are convergence points and sites at which many of these biological changes occur, including altered lipid metabolism, endoplasmic reticulum (ER) stress and an increase in reactive oxygen species. Consequently, we hypothesize that there will be essential concomitant changes in these organelles that reflect tumorigenesis. It is essential to understand the biological mechanisms and cell signaling that sustain these adaptations and result in tumor progression. For example, PCa displays significant alterations in endosome biogenesis and trafficking, which correlates with tumor progression and malignancy(1–3). While not yet characterized in PCa, other cancer types are dependent on vesicular transport trough the cytoplasm and organelle contact site biology(4–6). Whilst many of these effects have been characterized at a molecular level much less is known about the impact on the cellular ultrastructure as defined by the morphology, distribution and relationship between organelles such as mitochondria and the endoplasmic reticulum.

The morphological adaptations that organelles and organelle contacts in prostate cancer cells undergo during tumor progression are not known. The majority of the ultrastructural imaging research done to map the alterations in human prostate tissue was done in the 60s and 70s. Since then, we have significantly progressed our understanding of the cell biology that is associated with pro-proliferative, metastatic and anti-apoptotic pathways. Therefore, we found it timely to use transmission electron microscopy to investigate the ultrastructure of PCa tumor tissue to identify structural adaptations of fundamental membrane trafficking pathways and membrane contact sites (MCS). MCS are tight contact sites between organelles often involving the endoplasmic reticulum (ER); ER-mitochondria, ER-Golgi, ER-endosomes, lipid droplet-mitochondria(7–9). The ER is fundamental to the cell’s response to metabolic changes and response to stress. It maintains nutrient homeostasis, protein synthesis and secretion, glucose homeostasis, calcium signaling, lipid synthesis and lipid droplet biogenesis. The ER has evolved multiple pathways to adapt to stress and metabolic changes which include activation of the unfolded protein response (UPR), expanding the ER volume, sensing cholesterol concentrations, and remodeling its contact network with other organelles(10,11).

In this study we have focused on assessing changes in the ER morphology and mitochondria-associated ER contact sites in prostate epithelial cells in their native tissue environment and correlated the data with PCa histopathology. We hypothesized that membrane contact sites between mitochondria and ER would be enhanced in PCa to support cancer metabolism and inhibit apoptosis. From hereon we will call these contact sites mitochondria-associated membranes (MAMs). These contact sites form structural bridges between organelles and facilitate transfer of lipids, ions and signaling molecules and are more abundant in several metabolic and neurodegenerative disorders (10,12,13). These MCS are platforms for signaling that regulate cell survival and tumorigenesis, which include both oncogenes and tumor suppressors (8,14–17). Thus, despite being well established membrane structures with an important role in progression of breast and ovarian cancer they have not been characterized in PCa to date, nor has their androgenic regulation (6,18). Due to the small size of MAMs, which are defined as ER and mitochondria membranes at a distance of <25 nm apart, the only direct method with sufficient resolution for evaluating them is transmission electron microscopy (TEM).

The purpose of this study is to evaluate tissue samples from clinical prostatectomies by high resolution TEM to identify morphological changes in PCa epithelial cells from different disease grades. Moreover, we will examine androgen-dependent changes in these parameters using *ex vivo* tumor tissue culture.

## Materials and Methods

### Tissue samples

Fresh prostate tissue samples were obtained through the Australian Prostate Cancer BioResource from patients undergoing radical prostatectomy. None of the patients had received radiotherapy or androgen-deprivation therapy prior to surgery. A summary of the pathology reports from the patient samples are shown in **Table 1**. The use of the samples for research were approved by the Human Research Ethics Committee of the University of Adelaide (H-2012-016). Benign and malignant areas in fresh human prostate tissue were identified post-surgery from prostatectomies. Normal tissue adjacent to tumor tissue was used as a normal control. Following macro-dissection, the tissue was further processed for transmission electron microscopy (TEM), paraffin embedding and immunohistochemistry or *ex vivo* tissue culture.

**Table 1.** Pathology reports of PCa tissue samples used in this study.

| Patient Number | Surgical Biopsy Report | Secondary Pathology Report | P stage | TEM sample Pathology Report | AMACR stain | PIN4 Basal Cell stain |
| --- | --- | --- | --- | --- | --- | --- |
| 33169L | GS 3+3 | GS 6 (3+3) | PT2C | GS 6 | 90% | - |
| 33136L | GS 3+4 | GS 7 (3+4) | PT3A | GS 6 | 50% | +/- |
| 33149L | GS 3+4 | GS 7 (3+4) | PT2 | GS 6 | 10% | + |
| 33162L | GS 4+3 | GS 7 (4+3) | PT3A | GS 7 (4+3) | 80% | - |
| 33699R | GS 4+3 | GS 7 (4+3) | PT3A |  | 50% |  |
| 33162L | GS 4+5 | GS 9 (4+5+3) | PT3B |  |  |  |

The samples were divided into three groups based on the surgical biopsy and secondary pathology reports; “normal” tumor adjacent tissue, low grade tissue with Gleason scores 3+3 and 3+4, and an intermediate grade group for Gleason score 4+3 and 4+5. The grading of the tissue was further validated in a third independent analysis for 4 samples in sections neighboring those of the ultrathin sections used for TEM analysis (**Table 1** – labelled TEM sample pathology report).

### Patient-Derived Explant Culture

Tissue was dissected into 1 mm^3^ cubes and cultured in triplicate on gelatin sponges soaked in media in a humidified incubator with 5% CO_2_ as previously described(19,20). Tissue samples from adjacent areas were cultured *ex vivo* either in the presence of the enzalutamide (10µM), perhexiline (10µM) or DMSO for 72 hours.

### Transmission Electron Microscopy

Tissue samples (∼1µm^3^) were fixed in 1.25% glutaraldehyde, 4% paraformaldehyde in PBS, 4% sucrose (pH7.2), post-fixed in 1% osmium tetraoxide for an hour and dehydrated in a graded series of ethanol. The samples were incubated in propylene oxide for a final dehydration step followed by a 1:1 epoxy resin:propylene oxide incubation overnight. The samples were incubated in pure epoxy resin overnight (EM912 from EMS Science or Durcupan from Fluka) and cured at 50°C for 48 hours. Semithin (1µm) and ultrathin sections (70 nm) were prepared on a Leica Ultracut 7 ultramicrotome. Semithin sections were stained with Toluidine Blue and areas containing glandular structures were identified and subsequently trimmed for ultrathin sectioning. Ultrathin sections were collected on copper-iridium slot grids (EMS Science), stained with 2% uranyl acetate in water (Agar Scientific) and Reynold’s lead citrate (Fluka).

Electron micrographs were collected at 80kV in a Tecnai 12 and a Jeol 1200 transmission electron microscope. High resolution image analysis (15-20,000 x magnification) was performed to quantify mitochondria-ER contacts in epithelial cells.

### Image Analysis

For membrane contact sites 20 images per sample were analyzed (15-20kx). Each image contained a minimum of 4 mitochondria. The length of the mitochondria outer membrane and length of membrane contact with the ER was recorded. A mitochondria-ER membrane contact site was defined as being a minimum of 50 nm in length. A MAM, <25nm distance between ER and mitochondria and a loose contact, WAM, between rough ER and mitochondria ranging 25-80nm in distance.

### Statistical Analysis

Before performing statistical analysis, outliers were identified with a ROUT test and excluded. The Student’s t-test was used for comparison of groups using GraphPad Prism version 9.3.1 for macOS. P-values <0.05 were considered statistically significant. Data are expressed as mean ± standard error of the mean.

## Results

### Evaluation of epithelial cellular morphology in prostate cancer tissue by TEM

The prostate glandular tissue consists of stromal and epithelial cells. The columnar secretory epithelial cells are facing the lumen of the gland, have a pale cytoplasm and round nuclei. Below is a layer of basal cells with oval shaped nuclei and dark cytoplasm. Occasional neuroendocrine cells are observed at the base of a gland. Prostate adenocarcinoma is characterized by a loss of basal cells and an expansion of luminal epithelial cells resulting in a loss of the lumen. The histological features of the glands are used to grade tumors with the Gleason score (21,22). PCa most commonly arises from the luminal epithelial cells of the gland, and we have therefore focused our studies on this cell type using transmission electron microscopy (TEM) of Gleason graded tumor tissue. To identify glands in the tissue samples at light microscopic level we used semithin sections (**Figure 1A**), prepared adjacent ultrathin sections TEM (**Figure 1B**), identified luminal epithelial cells and imaged at high resolution (**Figure 1C**). In particular we focused on the organelles and structures associated with the endomembrane compartment that recently have received attention for promoting or sustaining cancer cell transformation; namely the Golgi, ER, lipid droplets and lysosomes.

**Figure 1:**
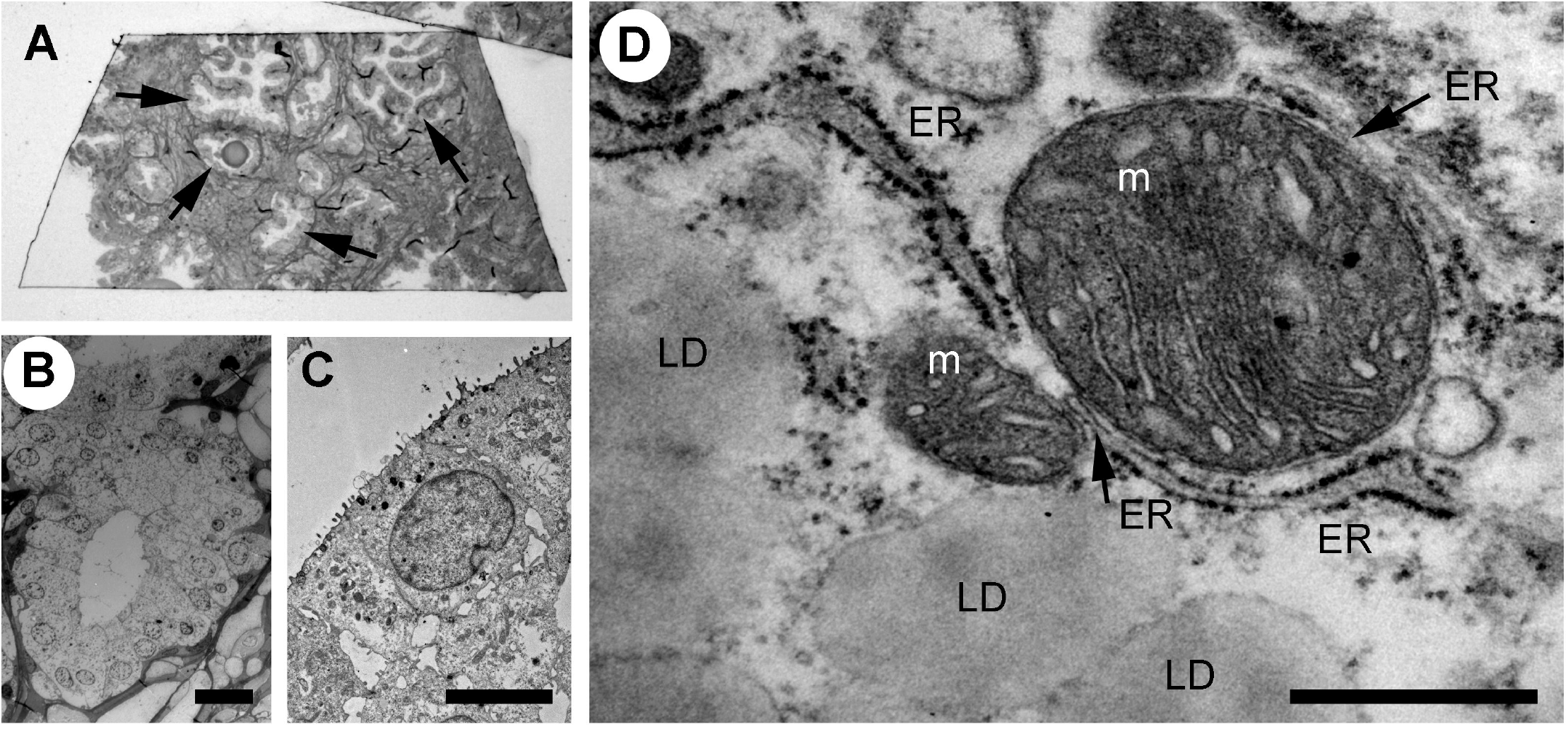
Analysis of the ultrastructure of secretory epithelial cells in human prostate tissue. **(A)** Glandular structures were identified in semi-thin sections of prostate tissue by light microscopy. Arrows point to glands. The sample was further trimmed down to regions containing glands and ultrathin section were prepared for TEM microscopy where the same structures were imaged at high resolution allowing secretory epithelial cells to be identified based on their morphology **(B)** Electron micrograph showing a gland at 100x magnification. Scale bar 20 µm. **(C)** Electron micrograph showing the secretory epithelia within a gland in the prostate. Scale bar, 5 µm. **(D)** Intracellular organelles in luminal epithelial cells were imaged at 8,000-20,000x magnification. Lipid droplet, LD; ER, endoplasmic reticulum; m, mitochondrion. Scale bar, 500 nm.

### Investigating potential Golgi adaptations in PCa cells

The Golgi apparatus is a central hub in the secretory pathway and is a fundamental site of complex N-glycosylation of lipids and proteins. Due to the specialized function of the prostate epithelia in secreting prostatic fluid, the Golgi apparatus is essential for production of secretory vesicles that contain proteins that promote sperm motility and facilitate semen liquefaction, including the prostatic-specific antigen (PSA). The Golgi has more recently been shown to have activities that are critical for cell death signaling, including localization of pro-apoptotic proteins and regulation of autophagy, stress sensor and mediator of malignant transformation(23). Recent evidence suggest that Golgi-mediated glycosylation is an important mediator of cell survival in prostate cancer and we therefore investigated if there were morphological changes in cancer cells compared to normal in prostate tissue.(24,25).

The Golgi is organized in stacks of flat cisternae that are held together by tethering proteins and cytoskeleton, which are further linked laterally in ribbons(26–28). The lipid composition of the Golgi membrane changes significantly from the *cis*-face to *trans*-face, where the composition more closely resembles that of the plasma membrane, with a higher concentration of cholesterol and sphingolipids. In the normal prostate epithelial cells, the Golgi had stacked cisternae with vesicles surrounding them (**Figure 2A**) and a ribbon organization (**Figure 2B**). A similar Golgi structure was observed in GS3+4 (**Figure 2C**) and GS4+3 samples, where Golgi stacks were closely aligned and slightly less dilated compared to the control (**Figure 2D**). In summary, no significant changes in the morphology of the Golgi were observed in our samples which may reflect that the luminal epithelial cells are secretory and inherently have a high capacity for protein processing.

**Figure 2:**
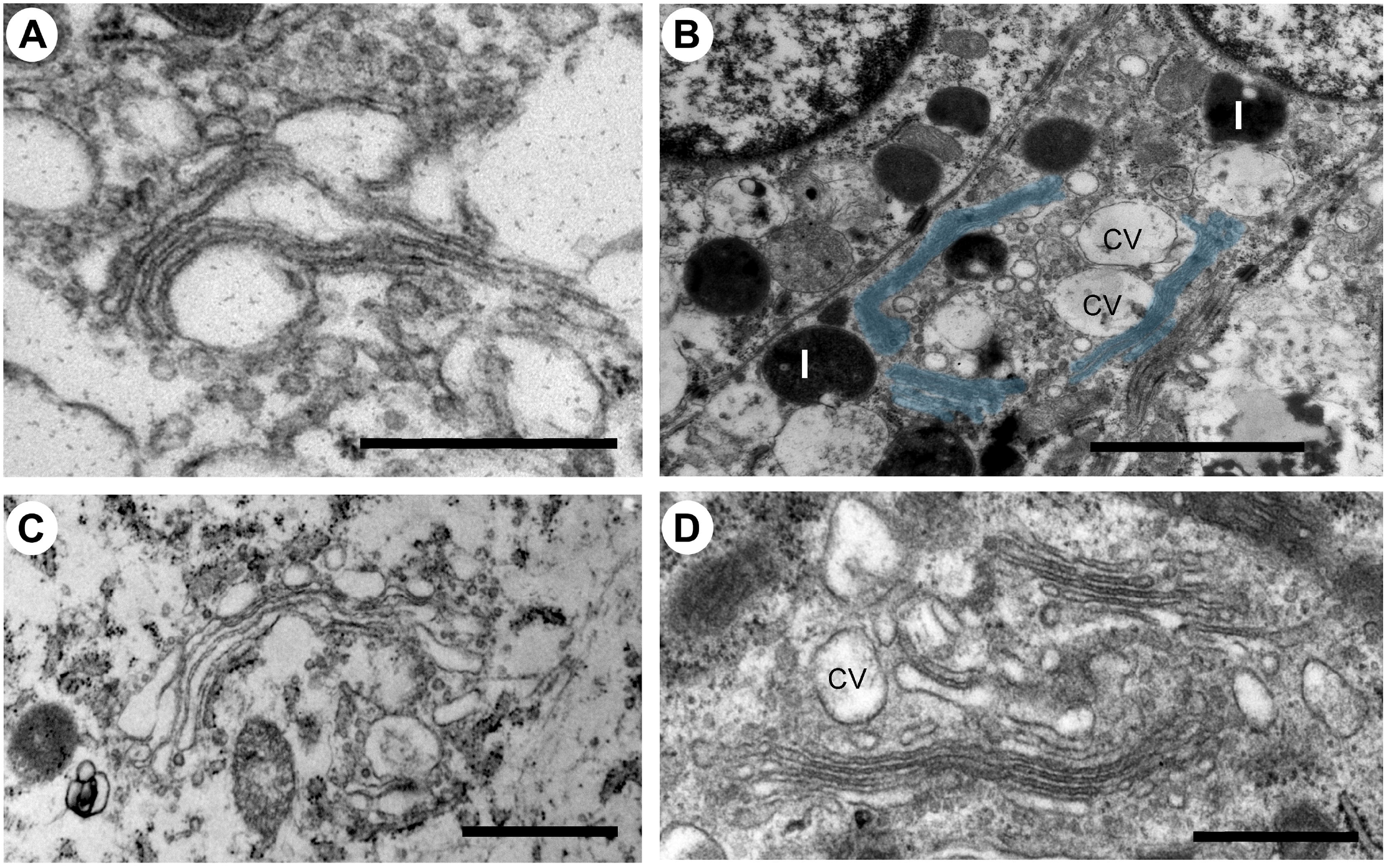
Ultrastructure of the Golgi apparatus in normal versus PCa tissue. **(A-D)** The Golgi apparatus in a secretory epithelial cell in a gland from a normal **(A)**, intermediate grade **(B, D)** and a low grade **(C)** prostate cancer tissue region had a normal appearance with stacked cisternae and a classic ribbon structure. The Golgi ribbon, pseudo colored in blue, measured up to 6 µm in length **(B)**. Scale bars, 500 nm (A, D), 2µm (B) and 1µm (C).

### Endoplasmic reticulum and lipid droplets in prostate epithelial cells

The endoplasmic reticulum (ER) is the largest organelle in the cell and is functionally involved in regulation of calcium signaling, protein synthesis, lipid biosynthesis and glucose metabolism(10). The ER is very adaptable and changes its morphology in response to the specialized function of the cell and to its metabolic status(29). The ER is formed by a network of membranes that is divided into three domains; the nuclear envelope, the perinuclear region consisting of ER sheets with ribosomes (rough ER) and the peripheral ER which a tubular network architecture (smooth ER). The relative ratio of rough versus smooth ER corresponds to the specialized function of a cell. Cell types that both secrete proteins and synthesize lipids, for example prostate epithelia cells, have an equal distribution of ER sheets and tubules (10,30). When the ER senses stress, a change in energy demand or the cellular metabolism, an adaptive change in morphology of the ER and its membrane interaction interfaces with other organelles will occur to maintain cellular homeostasis. Thus, an increase in the tubular ER promotes lipid metabolism and formation of lipid droplets (31,32). We next investigated the ER architecture in the prostate epithelia. We found a mixture of ER sheets and tubules in the different grade groups reflecting its specialization in protein synthesis/secretion and steroid synthesis (**Figure 3A, B**). In intermediate-high grade tissue ER whorls were observed that in some cases associated with lipid droplets **(Figure 3C)**. These structures have been shown to form due to high levels of HMG-CoA reductase, a key enzyme in the cholesterol biosynthesis pathway, central to the lipid reprogramming that occurs during PCa progression (33,34).

**Figure 3:**
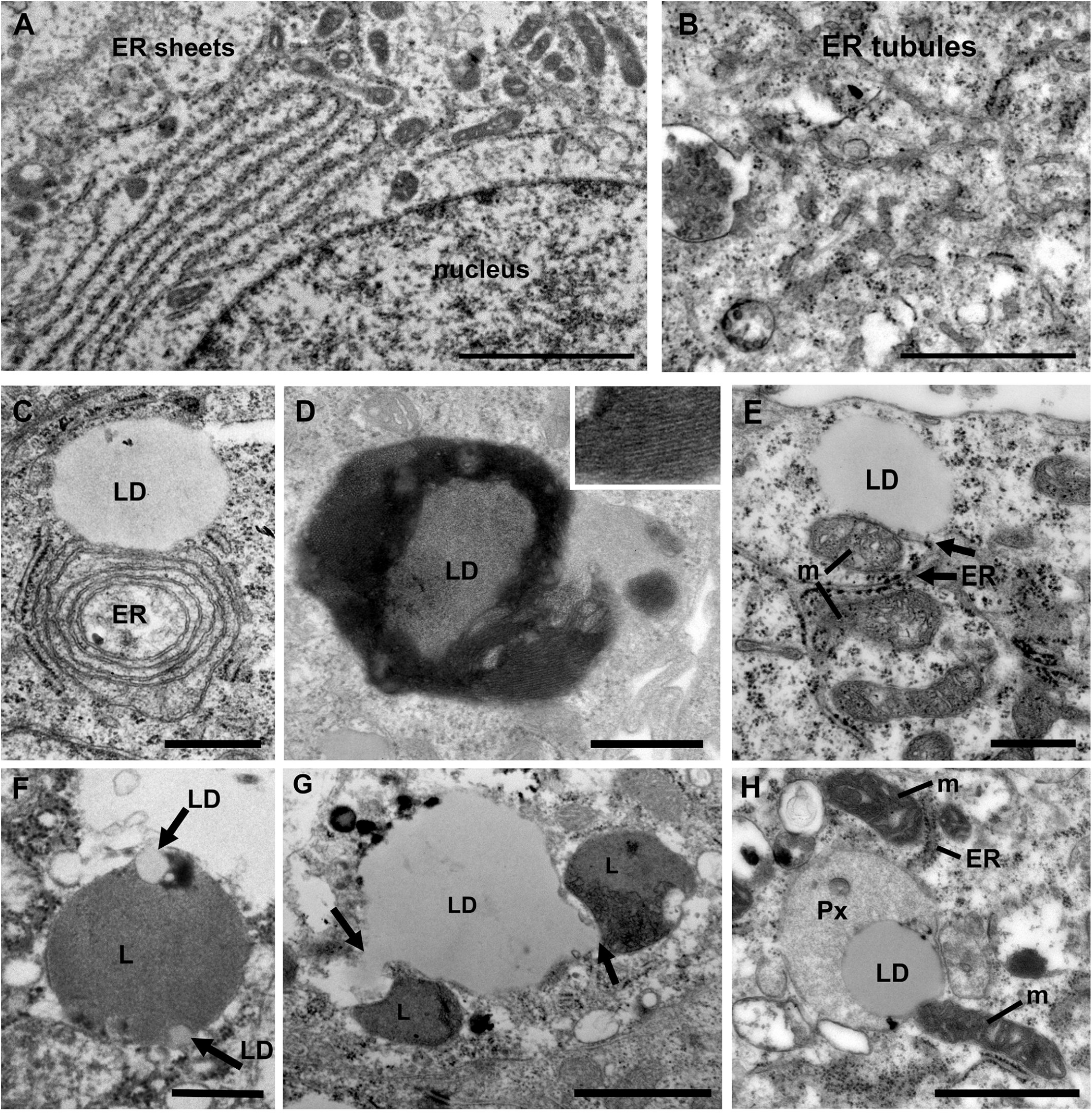
Ultrastructure of organelles and their membrane contacts in PCa tissue. **(A)** Electron micrograph illustrating the presence of ER sheets close to the nucleus (n) of secretory prostate epithelial cells. Scale bar, 2 µm. **(B)** ER tubules form a network throughout the cytoplasm in intermediate grade PCa tissue. Scale bar, 500 nm. **(C)** In intermediate grade PCa tissue lipid droplets were often found in contact with the ER. In this image an ER whorl is forming a membrane contact site with a lipid droplet (LD). **(D)** An electron micrograph from intermediate grade PCa tissue showing a lipid droplet with striations indicating a liquid-phase separation of lipids. Scale bar, 500 nm. **(E)** Electron micrograph showing membrane contact sites between a lipid droplet, mitochondria, peroxisome (Px) and ER in an intermediate grade PCa tissue. Scale bar, 1000 nm. **(F)** A micrograph showing an example of a lysosome engulfing small lipid droplets (arrows) in normal tissue. Scale bar, 500 nm. **(G)** Electron micrograph showing two lysosomes (arrows) engaged in lipophagy at a large lipid droplet (LD) in PCa tissue of intermediate grade. Scale bar, 1µm. (**H**) Electron micrograph illustrating membrane contact sites between lipid droplets, mitochondria (m) and (ER) in a luminal epithelial cell from a PCa tissue of intermediate grade. Scale bar, 500 nm.

Lipid droplets are organelles surrounded by a monolayer of lipid membrane that originate from the ER. In PCa cell lines treated with androgen the number of lipid droplets identified by Oil Red O staining is markedly upregulated (35). PCa tumor sections show an increase of lipid droplet staining compared to normal tissue (36). We observed lipid droplets attached to the ER (**Figure 3C**) and unattached (**Figure 3D**). Some lipid droplets in intermediate-to-high grade tissue showed evidence of storage of lipids in a phase separated fashion, known as smectic liquid-crystalline form (**Figure 3D**). This occurs when triglycerides stored in LDs are mobilized and the remaining cholesteryl ester enters a crystalline state which can be observed as a tight striation, shown in the inset. This is observed in cells undergoing metabolic stress along with an increase in LD-mitochondria membrane contact sites and reflects metabolic adaptations (37). Here we observed frequent contacts between lipid droplets and the ER (**Figure 3E**). However, no significant differences between Gleason score groups were observed (data not shown). Three-way organelle contacts between lipid droplets, mitochondria and ER were occasionally observed (**Figure 3E**).

An excess of lipids may lead to lipotoxicity and oxidative stress, and adaptive mechanisms in PCa to mitigate against such stress are therefore important to understand, particularly since an upregulation of lipid synthesis is characteristic of the disease(38). There are two key mechanisms that aid the cell in removing excess lipids; storage in lipid droplets and degradation of by lipophagy, a form of selective autophagy that specifically act on lipid droplets. The role of lipophagy in PCa is not fully understood but is proposed to associate with disease aggressiveness (36). Its role in other cancer types and metabolic disorders has made it an emerging biology of interest that requires further investigation in PCa (39). Here we observed for the first-time direct evidence of microlipophagy in luminal epithelial cells in normal and prostate cancer tissue (**Figure 3F, G)**. Microlipophagy is characterized by a direct contact between lysosomes and a lipid droplet. In normal tissue macrolipophagy typically occurred with small lipid droplets that were engulfed by a lysosome (**Figure 3F**). In PCa tissue, where large lipid droplets (>1µm) are more common, lysosomes were observed nibbling on and pulling out 150 nm wide tubes/vesicles from the lipid droplet (**Figure 3G**). In addition to mitochondria and lysosomes, lipid droplets were also found in direct contact with peroxisomes in PCa cells (**Figure 3H**). Peroxisomes play an important role in lipid metabolism and the activity of peroxisomes is upregulated in PCa (40,41). Here the lipid droplet was associated both with a peroxisome and a mitochondrion, and the mitochondria was in addition in contact with the ER. In summary, the use of high-resolution imaging of prostate luminal epithelial cells in tissue have demonstrated that membrane contacts between organelles exist and the organelles can engage in multiple contacts simultaneous, which may promote energy production and cell growth.

### Quantification of ER membrane contact sites with mitochondria in PCa tissue

Next, we investigated membrane contact sites between mitochondria and ER using a morphometric analysis of EM images. Mitochondria form two types of ER contact sites, MAM, that are juxtaposed to smooth ER and wrappER-associated mitochondria (WAM) that contact the rough ER. WAM contact sites regulate the biogenesis if very-low-density-lipoproteins (VLDL) in the liver and whole-body lipid homeostasis (42). MAMs are important structures for regulation of intracellular lipid metabolism, lipid synthesis, calcium signaling, mitochondrial function and apoptosis(5,6,10). Based on their function MAMs have been proposed to regulate cancer metabolism and fate. Their number, length and contact thickness have important impacts on their biological functions. Here we investigated the abundance of MAMs in prostate epithelial cells in their tissue environment. MAMs were defined as a membrane contact site between mitochondria and smooth ER that were within 25 nm. Electron micrographs show the contact sites in normal prostate (**Figure 4A**), low grade PCa tissue (**Figure 4B**) and intermediate-to-high grade PCa tissue. The abundance of MAMs was quantified as percentage of mitochondria that associated with ER membrane from three clinical samples per group (**Figure 4D**). A significant increase in MAMs was observed in epithelia from tissue in the intermediate-to-high grade group compared to normal tissue. In this group a mean of 71% of mitochondria were associated with ER membrane with an average of 81 mitochondria analyzed per clinical sample (**Figure 4D**). The MAM contact sites were also significantly longer compared to those in normal tissue in both the low and intermediate-to-high grade groups (**Figure 4E**). It is worth noting that the quantification does not reflect the total length of contact sites as single sections were analyzed in this experiment. This is the first analysis of MAMs in human prostate tissue to date. In addition to the MAMs we also quantified the abundance of mitochondria-rough-ER contacts (WAM) that have been described in detail by the Pellegrini group (42,43) (**Figure 4C**). We found a small but significant increase of these inter-organellar contacts and an increase in length that correlated with PCa progression to intermediate-to-high grade (**Figure 4F, G**). In summary, we have described a novel positive correlation between MAMs and PCa progression in luminal epithelial cells in their tissue environment.

**Figure 4:**
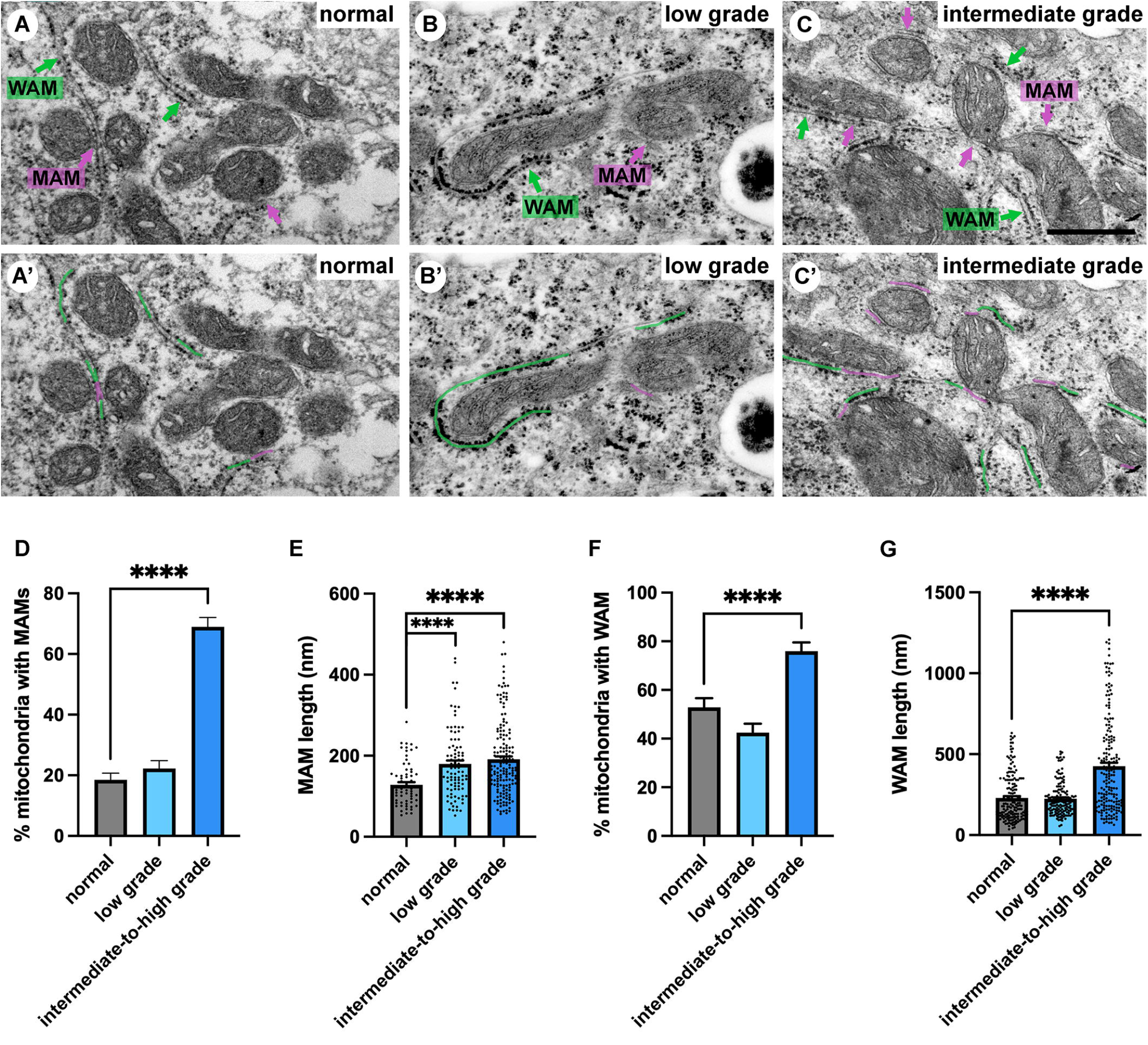
Ultrastructure of ER-mitochondria contacts in PCa tissue. **(A-C)** Ultrastructure of MAMs (magenta pseudo color) and wrappER-associated mitochondria (WAM) (green pseudo color) in luminal epithelial cells of normal, low and intermediate grade prostate cancer. Scale bar, 500nm. **(D, E)** Bar graphs showing the quantification of MAMs from normal (N=3), low grade (N=3) and intermediate-to-high grade (N=3) using 20 images per sample and >70 MAMs per group for length measurement. **(F, G)** Bar graphs showing the quantification of abundance and length of WAM contacts in the samples described in D. The length of WAM contacts were measured in a minimum of 150 mitochondria per group. All data are represented as mean ± sem; unpaired Student’s t-test. ****p<0.0001.

### Investigation of androgen-dependence of MAMs in prostate cancer

Gleason grading is an important predictor of PCa outcome. A tumor with a Gleason grade of 4+3 has a three-fold higher risk of progressing to lethal PCa compared to a Gleason grade 3+4 and is also predictive of later metastatic disease (44,45). The androgen receptor (AR) and its transcriptional activity is upregulated in PCa, resulting in over-expression of AR-target genes such as the lipid producing enzyme fatty acid synthase and a suite of other key lipid metabolic genes (46–48). Prostate cancer progression is dependent on androgen signaling though the AR, and consequently the first-line treatment strategy for locally advanced or metastatic disease is androgen-deprivation therapy (ADT). Due to cellular adaptations the cancer inevitabley develops therapy resistance to ADT, and the majority of patients are diagnosed with incurable castrate-resistance prostate cancer (CRPC) within 2-3 years. At this stage a standard of care treatment is the androgen receptor inhibitor enzalutamide that inhibits nuclear translocation, DNA binding and transcription of AR-target genes(49). Here we used enzalutamide-treated patient-derived explant tissues to investigate the impact of AR signaling on MAMs in intermediate-to-high graded tumor tissue (Gleason grade 4+3 and 4+5).

Patient-derived explant tissue was treated with enzalutamide for 72 hours *ex vivo* and processed for transmission electron microscopy analysis. The vehicle control-treated tissue (DMSO) had an ultrastructural morphology that was similar to control tissue that did not undergo *ex vivo* culture (**Figure 5A**). Samples treated with enzalutamide had a significantly reduced abundance of MAMs (**Figure 5B, D**). The number of wrappER-associated mitochondria contacts were not affected by enzalutamide treatment (data not shown). We quantified the length of 275 MAM contact sites from three independent patient-derived explant samples, which demonstrated a significant reduction in size of membrane contact site after enzalutamide treatment compared to the DMSO control (**Figure 5E**). As a control for our enzalutamide treatment we used samples treated with the drug perhexiline (**Figure 5C, F, G**). Perhexiline is a CPT-1 inhibitor and is used clinically to inhibit degradation of lipids for energy production by preventing the mitochondrial uptake of long-chain fatty acids. CPT-1 is an enzyme that is localized to the outer mitochondrial membrane. Treatment with perhexiline did not produce any significant changes to the abundance or length of MAMs. In agreement with studies showing an effect of perhexiline on cell proliferation and accumulation of autophagy markers we observed a large number of autophagic vacuoles in Perhexiline-treated samples, which was not observed in control cells thus providing evidence of the drug activity (**Figure 5H**; (50)). In summary, we have shown that MAMs are regulated by androgen signaling in prostate cells in their tissue environment. Future work will be important to elucidate the molecular mechanisms.

**Figure 5:**
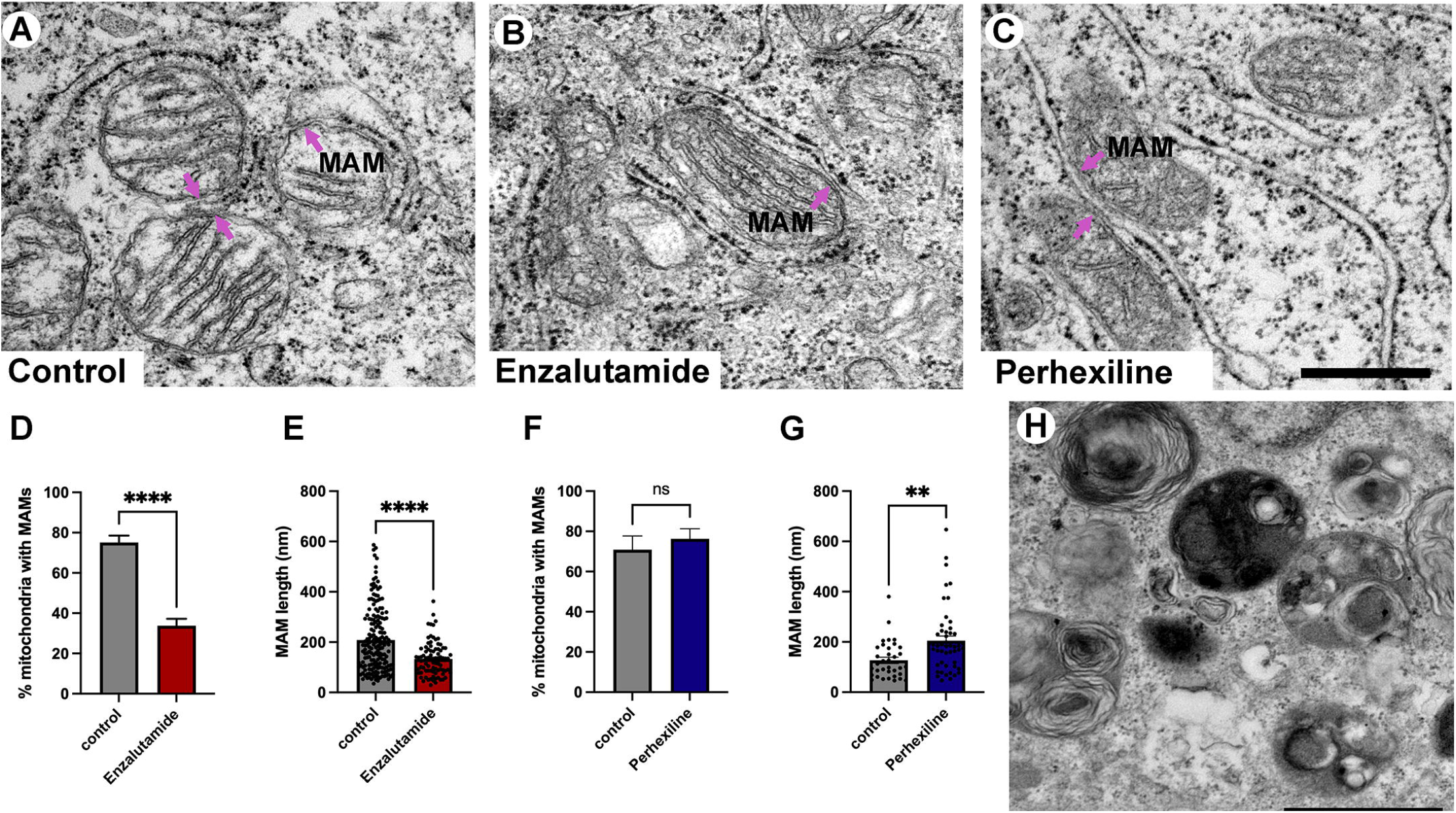
Quantification of MAMs from patient-derived tissue explants treated with the anti-androgen enzalutamide. **(A-C)** Electron micrographs of MAMs in DMSO-treated **(A)**, enzalutamide-treated **(B)** and Perhexiline-treated **(C)** explant tissue sections from PCa tissue with an intermediate-to-high Gleason score. Magenta arrows point to MAMs. Scale bar, 500 nm. **(D-E)** Analysis of enzalutamide-treated patient-derived tissue explant samples demonstrate a negative regulation of abundance and length of MAMs in three independent patient-derived tissue samples compared to paired DMSO-treated control samples. **(D)** Bar graph showing the percentage of MAM contacts among the total number of mitochondria. N=3, n≥20 images per sample. **(E)** Quantification of MAM length from electron micrographs (n=20 images) showed a significant difference between paired control-treated (DMSO) and enzalutamide-treated patient-derived tissue (N=3 clinical samples). We quantified the length of 183 control and 92 enzalutamide-treated MAMs. **(F)** Quantification of MAMs in Perhexiline-treated (C) tissue compared to a DMSO control from one patient-derived sample. The analysis included representative images; n=16 control and 20 treated. **(G)** Analysis of the length of MAMs in paired explant tissue samples treated with perhexiline. We quantified the length of 34 DMSO- and 49 perhexiline-treated MAMs. **(H)** Autophagic vacuoles in a perhexiline-treated explant tissue sample. Scale bar, 1 µm. All data are represented as mean ± sem; unpaired t-test. ****p<0.0001, ***p<0.001, **p<0.01.

## Discussion

Membrane contact sites formed by the ER are essential for facilitating communication between organelles and transporting ions and lipids across membranes. The adaptability of the ER and its ability to change its shape and architecture quickly in response to stress and cellular metabolism makes it an important organelle in cancer cell transformation. Recent research has shown an important correlation between membrane contact site formation between the ER and mitochondria for a number of diseases, including neurodegenerative disorders, cancer and obesity. In this study we investigated the correlation between MAM contact sites and PCa progression according to the histological Gleason grade. A key driver of PCa progression is androgen-dependent signaling, which include ER stress response mechanisms. A significant increase in MAMs in intermediate-to-high grade tumors were observed. Furthermore, we demonstrated that inhibition of AR target genes by treatment with the AR inhibitor enzalutamide in explant tissue results in a reduction in MAMs. Our results support a model in which MAM sites in prostate epithelial cells are regulated by AR signaling and/or transcriptional alterations that promote formation of contact sites and contribute to a biology that facilitates cell survival and proliferation.

The contacts between ER and mitochondria are mediated by MAM resident proteins and tethers. Protein-protein interactions from the two juxtaposing organelles form interaction platforms where the membranes align and allow lipids and ions to transfer, and in addition provide a platform for tumor suppressors and oncogenes to signal. These contact sites are enriched in cholesterol and sphingolipids and this membrane lipid composition has important implications for recruitment and insertion of membrane-associated proteins (51,52). Altered expression of these proteins have been reported in breast cancer, a hormone-dependent cancer, and have been shown to regulate cancer progression(6,53,54). For example, the MAM enriched chaperone sigma-1 receptor (Sig-1R) is significantly overexpressed in metastatic breast cancer compared to normal tissue(53,54). At the MAM Sig-1R regulates calcium fluxes to the mitochondria and cell survival together with BiP/GRP78 and inositol 1,4,5-trisphosphate receptors (IP3Rs) (55). In addition to adapting to changes in nutrient availability MAMs also responds to cellular stress by activating the unfolded protein response (UPR) (13). In prostate cancer lipid metabolism, ER stress and upregulation of the UPR are important mechanisms in the progression of disease (56–59).

What is the mechanism for increased MAMs in PCa? There are three main options. It could result from increased expression of cellular factors that stabilize the MAMs. Transcriptional regulation of genes by androgens via AR in PCa is a well-studied area and has identified enhanced expression of genes that support molecular mechanisms that promote cell proliferation and tumorigenesis (60–62). However, it is also possible that these changes in MAMs numbers/abundance is due to metabolic drivers that regulate the ER tubules-to-sheet ratio since the MAMs increase in cells upon ER tubule induction in the liver (29,63). Tissue-specific adaptations are common for formation of membrane contact sites and further research is required to elucidate the tubule/sheet ratio in the context of PCa progression. We did not find any evidence of changes in ER tubules-to-sheet ratio in our samples but cannot rule out that changes occur during PCa progression that may impact on MAMs. Ultrastructural studies in rats have shown a correlation between androgen signaling and ER structure in the prostate, where after castration a significant remodeling of the ER with a loss of cisternae and an increase in whorls was observed (33,64). Finally, the saturation of phospholipids and sphingolipids is known to impact on the MAMs content, which can promote mitochondrial fission at these sites (10,65,66). This is interesting in the context of PCa lipid metabolism since significant changes have been observed in chain length and saturation of lipids(67).

Lipid metabolism is elevated in PCa cells and contributes substrates for intra-tumoral steroid hormone synthesis. An increase in lipid droplet abundance is an established part of the progression from a low to high Gleason grade disease and is important because lipid droplets bud from the ER (35). The metabolism of lipid droplets and generation of energy are mediated by mitochondria, lysosomes and peroxisomes. Work in hepatocytes have shown that microlipopahgy is an important mechanism for lipid droplet metabolism (68,69) along with organelle contacts with mitochondria(70). Here we have shown for the first time that prostate epithelial cells can mobilize lipids stored in lipid droplets by microlipophagy. Prostate cancer aggressiveness and progression is positively correlated with markers of macrolipophagy, autophagic degradation of lipid droplets, and has been suggested to contribute to treatment resistance thus making our observations on microlipophagy highly interesting for further investigations (36,71). Recent research has shown that mitochondrial metabolism of lipids is regulated by organelle interactions and MAM around mitochondria(70). Short- and medium-chain fatty acids are used by the cell to generate energy through mitochondrial β-oxidation, but this is not possible for very-long-chain fatty acids which instead are catabolized in peroxisomes to short-chain fatty acids that then can be further oxidized in mitochondria. In most cells the energy gain from very-long-chain fatty acids is small due to their low abundance(72). However, PCa have an upregulation of the enzymes required for elongating fatty acids and this is reflected in the lipid profile generated by lipidomics (73). Our observations of direct membrane contact sites between peroxisomes, lipid droplets and mitochondria are therefore interesting and may reflect a rewiring that reflects the energy demands of PCa cells similar to what has been observed in the liver(74).

According to our data presented here, targeting MAMs may disrupt a cellular adaptation of prostate cancer cells that contribute to resolving stress, calcium homeostasis and lipid homeostasis. Further mechanistic studies using cell lines and explant tissue are needed to gain a better understanding of the molecular components that are fundamental for these contact sites in prostate cancer cells and whether there are metabolic alterations that contribute to establishment of MAMs in addition to androgen-regulated gene expression. This will ultimately provide insights to cancer cell survival, proliferation and disease progression. To conclude, our data have characterized a novel AR-regulated biology that is associated with PC disease progression. MAMs are important cellular signaling platforms for apoptosis, managing lipid transfer into mitochondria and are sites of lipid droplet biogenesis. The MAM structure therefore has the potential to direct/control adaptations that occur during PCa development that are very important in promoting survival in cancer cells.

## Ethics Statement

The project was approved by the Human Research Ethics Committee of University of Adelaide (H-2012-016). Informed written consent was obtained from patients to use surplus tissue after diagnosis for research through the Australian Prostate BioResource under approval from the St Andrews Hospital Human Research Ethics Committee (#80).

## Author Contributions

EE and LMB designed the research study, wrote the manuscript and approved the submitted article. EE performed the experiments.

## Acknowledgements

Tissues for the patient-derived explants used in the study were collected with informed consent via the Australian Prostate Cancer BioResource and we thank the doctors, patients and health care professionals involved. We acknowledge expert technical assistance in the study from Swati Irani, Kayla Bremert and pathology input from Dr Jurgen Stahl. The authors are grateful for the expert assistance from the Image Core Unit at Adelaide University and Queen’s University Belfast.

